# Spatial variation in black-headed night monkey (*Aotus nigriceps*) vocalizations

**DOI:** 10.1101/688333

**Authors:** W. D. Helenbrook, N. A. Linck, M. A. Pardo, J. A. Suarez

**Affiliations:** State University of New York College of Environmental Science and Forestry (SUNY-ESF), 1 Forestry Drive, Syracuse, New York, United States; Tropical Conservation Fund, 760 Parkside Trl NW, Marietta, Georgia, USA; University of Minnesota Twin-Cities, St. Paul, Minnesota, USA; Cornell University, 215 Tower Road, Ithaca, New York, USA

**Keywords:** vocalizations, *Aotus nigriceps*, night monkey, Peru, acoustics

## Abstract

Quantitative acoustic analysis has been used to decipher individual differences, population structure, and taxonomic diversity in numerous primate species. We previously described three distinct call types in wild *Aotus nigriceps*, and now assess acoustic differences in two of these call types between social groups and spatially distinct populations. Acoustic parameters for both analyzed call types exhibited significant variability between groups. Similarly, geographically distant field sites were acoustically distinct from one another. Several groups also used a variation of a common call: a triplet Ch Ch instead of a duplicate. Other groups made use of ultrasonic frequencies which have not previously been reported in *Aotus*. Our results suggest that *Aotus nigriceps* exhibits substantial acoustic variability across sites that could potentially be useful for taxonomic classification, although additional geographically distant populations still need to be sampled. The possibility of individual signatures also exists and will require recording vocalizations from known individuals.

## Introduction

Primates use vocalizations to communicate about the presence of predators (Zuberbühler 2002; Schel et al. 2009), location of food sources (Slocombe and Zuberbühler 2005), nesting behavior, travel intentions and group cohesion (Boinski 1996), territorial defense (Raemaekers and Raemaekers 1985; Cowlishaw, 1992), mate assessment and pair bonding (Cowlishaw 1996; Geissmann and Orgeldinger 2000). Primate acoustic signals may also be used for kin recognition and can convey information on age, sex, body size, and rank (Salmi and Hammerschmidt 2014). The diversity of these vocal signals makes acoustics a convenient tool to explore differences among primates at various taxonomic levels.

Despite the importance of acoustic communication in primates, evidence of population level differences in primate vocalizations are relatively limited (Green 1975; Maeda and Masataka, 1987; Mitani et al., 1992; Fischer et al. 1998; Mitani et al. 1999; Delgrado 2007; Wich et al. 2008; de la Torre and Snowdon 2009; Wich et al. 2012). Acoustic variation between populations can exist for any of several reasons: divergence through cultural drift in species that learn their vocalizations (i.e. inaccurate copying transmitted vertically or horizontally), genetic drift following reproductive isolation, or local adaptation in response to sexual selection, habitat transmission properties, predation pressure, or social selection pressures (Yoktan et al. 2011).

Most evidence suggests that non-human primates are not vocal learners; however, several recent studies have found that primates can learn slight modifications to their vocalizations (Watson et al. 2015; Takahashi et al. 2017). Examining patterns of geographic variation in call structure can provide some evidence for or against the presence of vocal learning; for example, if there is a sharp acoustic divide between two spatially contiguous areas that show no evidence of genetic divergence (i.e. vocal dialects), this often suggests the presence of vocal learning.

Spatial variation in vocal characteristics within a species could be a valuable tool for addressing a variety of ecological questions. For example, primate calls can be used to distinguish species, populations, groups, and individuals (Table 1). If individuals could be distinguished from one another solely using quantitative acoustic analysis, population size could be estimated by combining acoustic analysis and line transect surveys (Terry et al. 2005; Marques et al. 2013; Kalan et al. 2015). Moreover, variation in specific call characteristics could be used to infer group membership, or be used for taxonomic classification, supplementing morphometric or genetic data.

**Table 1.** Evidence of primate acoustic variation at various taxonomic levels.

| Primate species (N) | Individual | Group | Population | Subspecies | Species | Call description | Study |
| --- | --- | --- | --- | --- | --- | --- | --- |
| *Cebidae, Callitrichinae* (28) | - | - | - | - | Yes | Long call | Garbino 2018 |
| *Galago* spp. (8) | - | - | - | - | Yes | Loud call | Zimmermann 1990 |
| *Galago alleni* | - | - | - | - | Yes | Loud call | Ambrose 2003 |
| *Galago crassicaudatus, G. garnettii* | - | - | - | - | Yes | Loud call | Masters 1991 |
| *Galago senegalensis, G. moholi* | - | - | - | - | Yes | 14 calls | Zimmermann 1988 |
| *Gorilla gorilla* | Yes | - | - | - | - | 8 calls | Salmi et al. 2014 |
| *Hylobates muelleri* | Yes | - | No | - | - | Great call | Clink et al. 2017 |
| *Hylobates muelleri* | Yes | - | Yes | - | - | Great call | Clink et al. 2018 |
| *Lepilemur* spp. (10) | - | - | Yes | - | - | Loud call | Méndez-Cárdenas et al. 2008 |
| *Microcebus* spp. (3) | - | - | - | - | Yes | Whistle, Purr, Chitter | Hending et al. 2017 |
| *Microcebus* spp. (3) | - | - | - | - | Yes | Advertisement call | Braune et al. 2008 |
| *Microcebus murinus* | - | - | Yes | - | - | Trill | Hafen et al. 1998 |
| *Tarsius* spp. (4) | - | - | - | - | Yes | Loud call (Duet call) | Nietsch 1999 |
| *Varecia variegata* | - | - | - | Yes | - | Loud call | Macedonia and Taylor 1985 |

Night monkeys, *Aotus* spp., are a useful model for investigating patterns of acoustic variation because nocturnal and forest-dwelling species tend to rely heavily on vocalizations to communicate with one another. We have previously reported on the vocal repertoire of wild *Aotus nigriceps*, describing three calls: the Squeak, Ch Ch, and Long Trill (Helenbrook et al. 2018). In this study we focus on quantitatively comparing acoustic variation of two of these calls between groups and distant populations.

## Methods

Eleven *Aotus nigriceps* groups were sampled (Fig.1): eight at the Villa Carmen Biological Station in Pilcopata, Peru (12°53’39"S, 71°24’16"W), and three at CREES - the Manu Learning Center, on the edge of Manu National Park (12°47′22″S 71°23′32″W). The two field sites are separated by a low mountain range (~1143m) and are just over 10 km apart at their nearest borders. Villa Carmen has a long history of development, ecotourism and agriculture. The groups sampled near the station lived in secondary forest, often dominated by bamboo or cane, whereas groups sampled at CREES inhabited recovering clear-cut to primary rainforest where bamboo and cane were largely absent. Research groups of 3-8 observers went into the field from 5:30-7:30am and 5:30-7:30pm for a total of 28 days at Villa Carmen and nine days at CREES to collect acoustic data, times when *A. nigriceps* groups are known to be active near their nesting sites. Several recordings also took place during the day as part of a separate behavioral study.

**Fig. 1.**
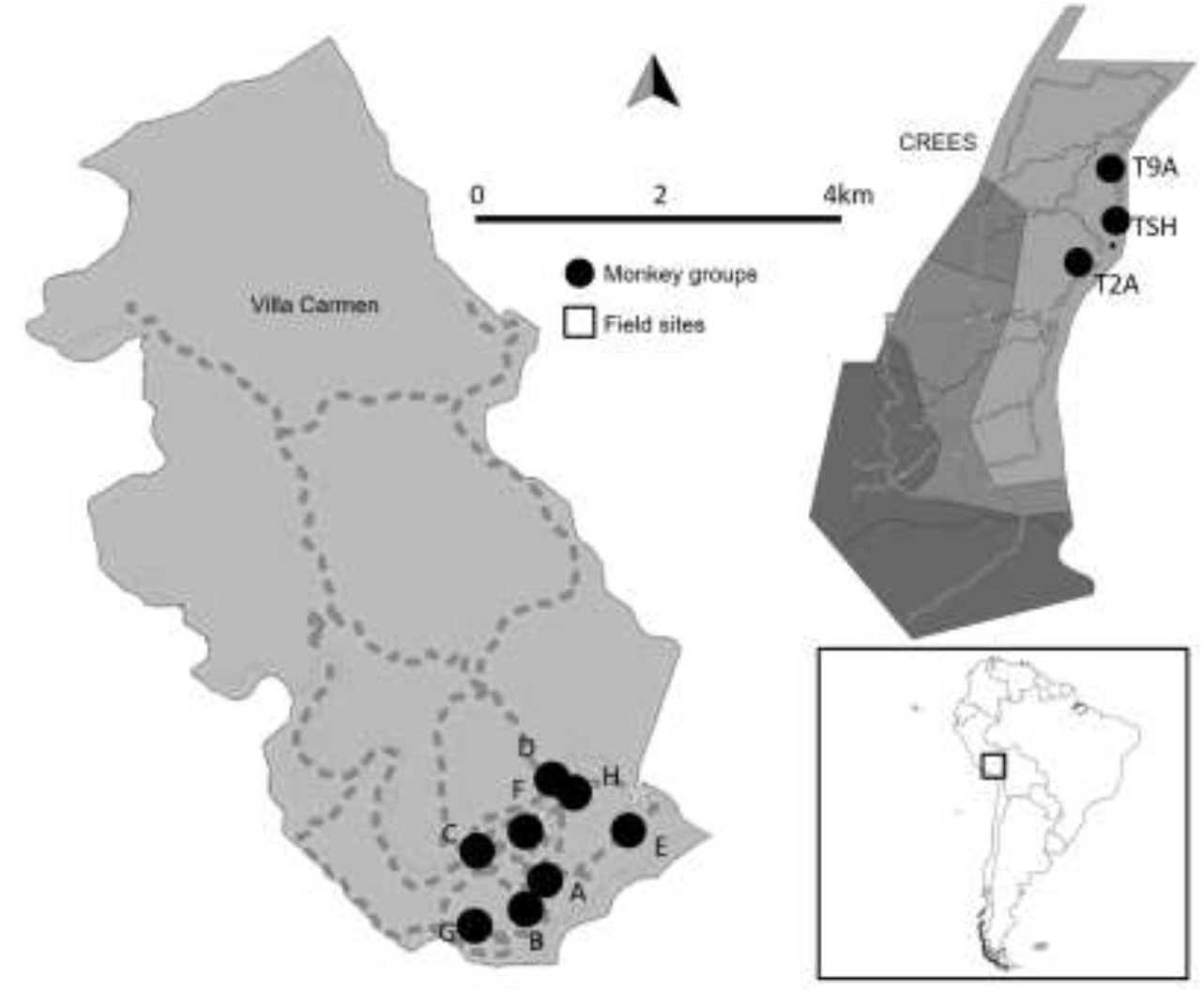
Field research conducted at Villa Carmen Biological Station (eight black-headed night monkey groups) and CREES – Manu Learning Centre (three groups), in southeastern Peru. A mountain ridge separates the two field stations (10.1km at nearest points)

A Zoom H1 Handy Recorder was coupled with a RØDE NTG-2 condenser shotgun microphone and shoe shockmount on a micro boompole at a distance varying from 2-25m. Digital recordings were made at 48 kHz sampling frequency with 16 or 24-bit amplitude resolution. Acoustic analysis was conducted using Raven Pro 1.5 sound analysis software (Cornell Lab of Ornithology Bioacoustics Research Program, Ithaca, New York). Calls were digitized and measured spectrographically (DFT size 512, time resolution 3.1 ms, Hann window with 50% overlap). Twenty-four acoustic parameters were measured for each call (Table 2).

**Table 2.** Name and description of acoustic parameters measured. Not all parameters were included in discriminant function analysis because of redundancy.

| Parameters | Description |
| --- | --- |
| Low frequency (Hz) | The lower frequency bound of the selection |
| High frequency (Hz) | The upper frequency bound of the selection |
| Bandwidth 90% | The difference between the 5% and 95% frequencies |
| Energy (dB) | The total energy within the selection bounds |
| Dur90% | The difference between 5% and 95% times |
| Delta frequency (Hz) | The difference between the upper and lower frequency limits of the selection |
| Peak frequency | The frequency at which max power occurs within the selection |
| Delta time (s) | The difference between the begin and end time for the selection |
| Center frequency (Hz) | The frequency that divides the selection into two frequency intervals of equal energy |
| Q1 frequency (Hz) | The frequency that divides the selection into two frequency intervals containing 25% and 75% of the energy in the selection |
| Q3 frequency (Hz) | The frequency that divides the selection into two frequency intervals containing 75% and 25% of the energy in the selection |
| Max power | The maximum power in the selection. |
| Frequency 5% | The frequency that divides the selection into two frequency intervals containing 5% and 95% of the energy in the selection |
| Frequency 95% | The frequency that divides the selection into two frequency intervals containing 95% and 5% of the energy in the selection |
| Center time | The point in time at which the selection is divided into two time intervals of equal energy |
| Q1 time | The point in time that divides the selection into two time intervals containing 25% and 75% of the energy in the selection |
| Q3 time | The point in time that divides the selection into two time intervals containing 75% and 25% of the energy in the selection |
| IQR duration (s) | The difference between the 1<sup>st</sup> and 3<sup>rd</sup> quartile times |
| Time 5% | The point in time that divides the selection into two time intervals containing 5% and 95% of the energy in the selection |
| Time 95% | The point in time that divides the selection into two time intervals containing 95% and 5% of the energy in the selection |
| Max time | The first time in the selection at which a spectrogram point with power equal to max power occurs |
| Quantitative acoustic characteristics analyzed (Raven). |  |

Inter-group differences were analyzed using non-parametric Kruskal-Wallis tests coupled with post-hoc multiple comparisons of mean ranks tests with a Bonferroni correction. A Mann-Whitney *U* test was used to assess site differences between Villa Carmen and CREES. Stepwise discriminant function analysis was used to explore acoustic parameters that could be used to classify social groups. For model selection, a stepwise forward method was used (statistic, Wilk’s Lamda) with the criteria *F*_to enter_=3.84 and *F*_to remove_=2.71, and a tolerance level of <0.01 (STATISTICA). This process was repeated for both call types separately. Variables that failed a tolerance test where there was an almost exact linear relationship with other variables, did not enter the analysis. We used a 10-fold cross validation in which 90% of the calls were randomly chosen to calculate discriminant functions, while 10% was excluded for testing. Differences between observed and expected frequencies of duplicate versus triplicate Ch Ch calls was measured using Fisher’s Exact Test. All recordings were conducted non-invasively, minimized impact on behavior, and avoided excessive disturbance, and were therefore deemed exempt from the Institutional Animal Care and Use Committee approval. All applicable international, national, and/or institutional guidelines for the care and use of animals were followed.

## Results

Three vocalizations have been described in wild *Aotus nigriceps* populations: Squeak, Ch Ch, and Trill (Helenbrook et al. 2018). In this study, we analyzed acoustic variability for the two most common calls, the Squeak (N=1302) and the Ch Ch (N=556; Fig. 2). For Squeaks we only measured the dominant harmonic since it was consistently found across all sampled groups. The Trill was not used because of its rarity across most groups. At least ten calls were analzyed from each of seven night monkey groups, ranging in size from 2-5 individuals (Mean=3.7). The dominant harmonic of the Squeak ranged from a mean minimum frequency of 1591 Hz (Range: 74-3055 Hz; SD 470) to a mean maximum frequency of 2742 Hz (Range: 2010-4443; SD 252). The Ch Ch call ranged from a mean minimum frequency of 1698 Hz (Range: 44-9092; SD 1182) to a mean maximum frequency of 11636 Hz (Range: 3109-23726 Hz; SD 2538).

**Fig. 2.**
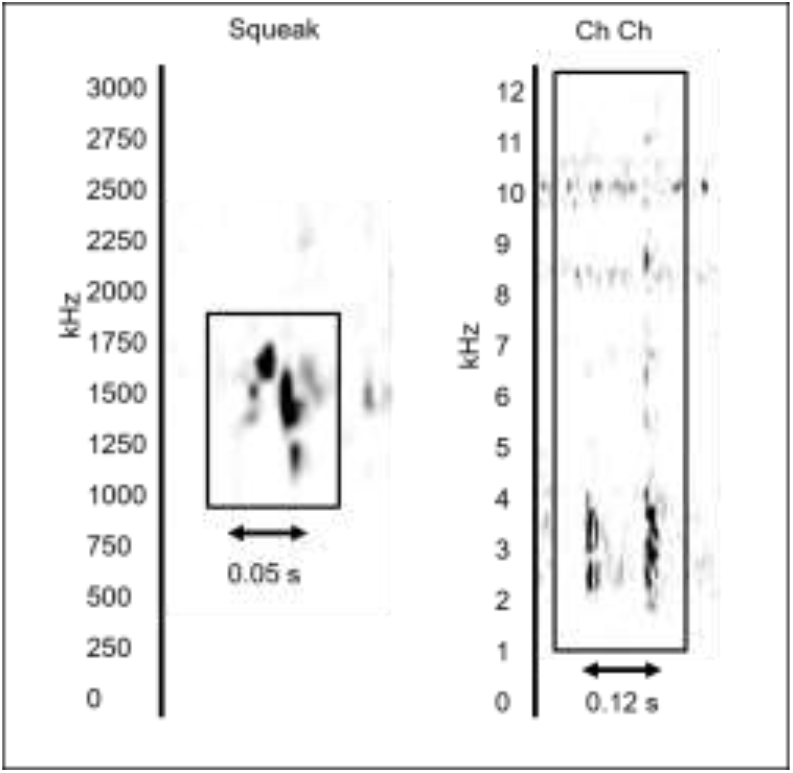
Spectrogram representing both calls quantitatively analyzed in this study: the Squeak (left) and the Ch Ch (right). Both calls have been previously described (Helenbrook et al. 2018). In this example, a duplicate Ch Ch is depicted. The upper range of the spectrogram is faint, partially a result of minimizing backgound noise in the 7-8 kHz from insects

Acoustic measurements varied significantly between groups (Fig. 3; Table 3). Differentiation between two or more monkey groups was found for all forty-eight independent vocal characteristics (2 call types x 24 measurements, p<0.001). Discriminant function analysis distinguished among groups for both call types: Squeak (Wilks’ Lambda=0.06, F(60,6706)=81.98, p<0.0000) and Ch Ch (Wilks’ Lambda=0.06, F(48,2592)=42.58, p<0.0000) (Table 4 and Fig. 4). Cumulative significant functions were able to explain 87.4% of variance among groups using only Squeak calls, and 87.8% of the variance among groups using only Ch Ch calls. Classification accuracy was similar for both the Squeak (87.4%) and the Ch Ch (76.4%). Duration (90%) and energy parameters for Squeak and Ch Ch, respectively, provided the greatest discriminatory power at the group level (Table 4).

**Table 3.** All *Aotus nigriceps* vocal measurements were found to have significant (p<0.001) inter-group differences. Post-hoc analysis was conducted using multiple comparisons of mean ranks with Bonferroni adjustment. All associations significant at p<0.05.

| Call Measurement | Group | Significant Associations (p value) |
| --- | --- | --- |
| *Ch Ch* |  |  |
| Dur90% | A | C(0.00); D(0.04) |
|  | B | C(0.00); D(0.01) |
|  | C | E(0.00); F(0.01); T2A(0.00) |
| High frequency | A | C(0.00); D(0.02); T2A(0.00) |
|  | B | C(0.00); D(0.04); T2A(0.00) |
|  | C | E(0.00); F(0.00); H(0.01) |
|  | D | E(0.01); F(0.01); T9A(0.05) |
|  | E | T2A(0.00) |
|  | F | T2A(0.00) |
| Low frequency | A | B(0.00); D(0.03); T2A(0.00) |
|  | B | C(0.00); D(0.00); E(0.00); T2A(0.00) |
|  | C | T2A(0.02) |
| Bandwidth 90% | A | B(0.00); E(0.00) |
|  | B | C(0.00); E(0.00); F(0.00); H(0.00); T2A(0.00); TSH(0.00) |
|  | E | T2A(0.00) |
|  | TSH | T9A(0.04) |
| *Squeak* |  |  |
| Dur90% | A | B(0.00); E(0.01); TSH(0.00) |
|  | B | C(0.02); E(0.00); T2A(0.00); TSH(0.00) |
|  | C | E(0.00); TSH(0.00) |
|  | E | H(0.00); T2A(0.00); TSH(0.00); T9A(0.05) |
|  | H | TSH(0.00) |
|  | T2A | TSH(0.00) |
| High frequency | A | C(0.03); E(0.00); H(0.00); TSH(0.02) |
|  | B | C(0.00); E(0.00); H(0.00); TSH(0.00) |
|  | C | E(0.00); H(0.00); T2A (0.00) |
|  | D | TSH(0.04) |
|  | E | H(0.00); T2A(0.00); TSH(0.00) |
|  | H | T2A(0.00); TSH(0.00) |
| Low frequency | A | B(0.00); E(0.00) |
|  | B | C(0.00); D(0.01); E(0.00); H(0.00); T2A(0.00); TSH(0.00); T9A(0.00) |
|  | C | T2A(0.00) |
|  | E | T2A(0.00) |
|  | H | T2A(0.00) |
| Bandwidth 90% | A | B(0.00); E(0.00); H(0.00); TSH(0.01) |
|  | B | C(0.00); E(0.00); H(0.00); T2A(0.00) |
|  | C | E(0.00); H(0.00); T2A(0.00) |
|  | E | T2A(0.00); TSH(0.00) |
|  | H | T2A(0.01); TSH(0.00) |

**Table 4.** Stepwise discriminant function analysis of all non-redundant measured acoustic parameters.

| Call Measurement | Wilks' Lambda | Partial Lambda | p value |
| --- | --- | --- | --- |
| *Squeak* |  |  |  |
| Bandwidth 90% | 0.07 | 0.85 | 0.00 |
| Energy | 0.06 | 0.92 | 0.00 |
| Dur90% | 0.07 | 0.82 | 0.00 |
| Frequency 95% | 0.06 | 0.88 | 0.00 |
| Max power | 0.06 | 0.89 | 0.00 |
| IQR duration | 0.06 | 0.96 | 0.00 |
| Center frequency | 0.06 | 0.98 | 0.00 |
| Q1 frequency | 0.06 | 0.93 | 0.00 |
| IQR bandwidth | 0.06 | 0.90 | 0.00 |
| Max time | 0.06 | 0.88 | 0.00 |
| *Ch Ch* |  |  |  |
| Energy | 0.17 | 0.33 | 0.00 |
| Dur90% | 0.06 | 0.92 | 0.00 |
| Bandwidth 90% | 0.05 | 0.94 | 0.00 |
| IQR duration | 0.06 | 0.96 | 0.00 |
| Frequency 95% | 0.06 | 0.89 | 0.00 |
| Center frequency | 0.09 | 0.65 | 0.00 |
| Delta frequency | 0.07 | 0.77 | 0.00 |
| Q1 frequency | 0.07 | 0.79 | 0.00 |

**Table 5.** Discriminant function analysis correct classification utilizing 10-fold cross validation at group level.

| *Squeak alone* |  |  |  |  |  |  |  |  |
| --- | --- | --- | --- | --- | --- | --- | --- | --- |
| Group | % Correct | A | B | C | E | H | T2A | TSH |
| A | 66.7 | **6** | 0 | 3 | 0 | 0 | 0 | 0 |
| B | 75.0 | 4 | **12** | 0 | 0 | 0 | 0 | 0 |
| C | 78.6 | 0 | 2 | **11** | 1 | 0 | 0 | 0 |
| E | 89.9 | 4 | 1 | 0 | **62** | 1 | 1 | 0 |
| H | 75.0 | 0 | 0 | 0 | 2 | **6** | 0 | 0 |
| T2A | 44.4 | 3 | 0 | 0 | 1 | 0 | **4** | 1 |
| TSH | 75.0 | 0 | 0 | 1 | 0 | 0 | 0 | **3** |

| *Ch Ch alone* |  |  |  |  |  |  |  |  |
| --- | --- | --- | --- | --- | --- | --- | --- | --- |
| Group | % Correct | A | B | C | E | F | H | T2A |
| A | 100 | **7** | 0 | 0 | 0 | 0 | 0 | 0 |
| B | 83.3 | 2 | **10** | 0 | 0 | 0 | 0 | 0 |
| C | 83.3 | 0 | 1 | **5** | 0 | 0 | 0 | 0 |
| E | 66.7 | 3 | 1 | 0 | **10** | 0 | 0 | 1 |
| F | 0 | 1 | 1 | 0 | 0 | **0** | 0 | 0 |
| H | 0 | 0 | 0 | 0 | 1 | 0 | **0** | 1 |
| T2A | 90.9 | 0 | 0 | 0 | 1 | 0 | 0 | **10** |

**Fig 3.**
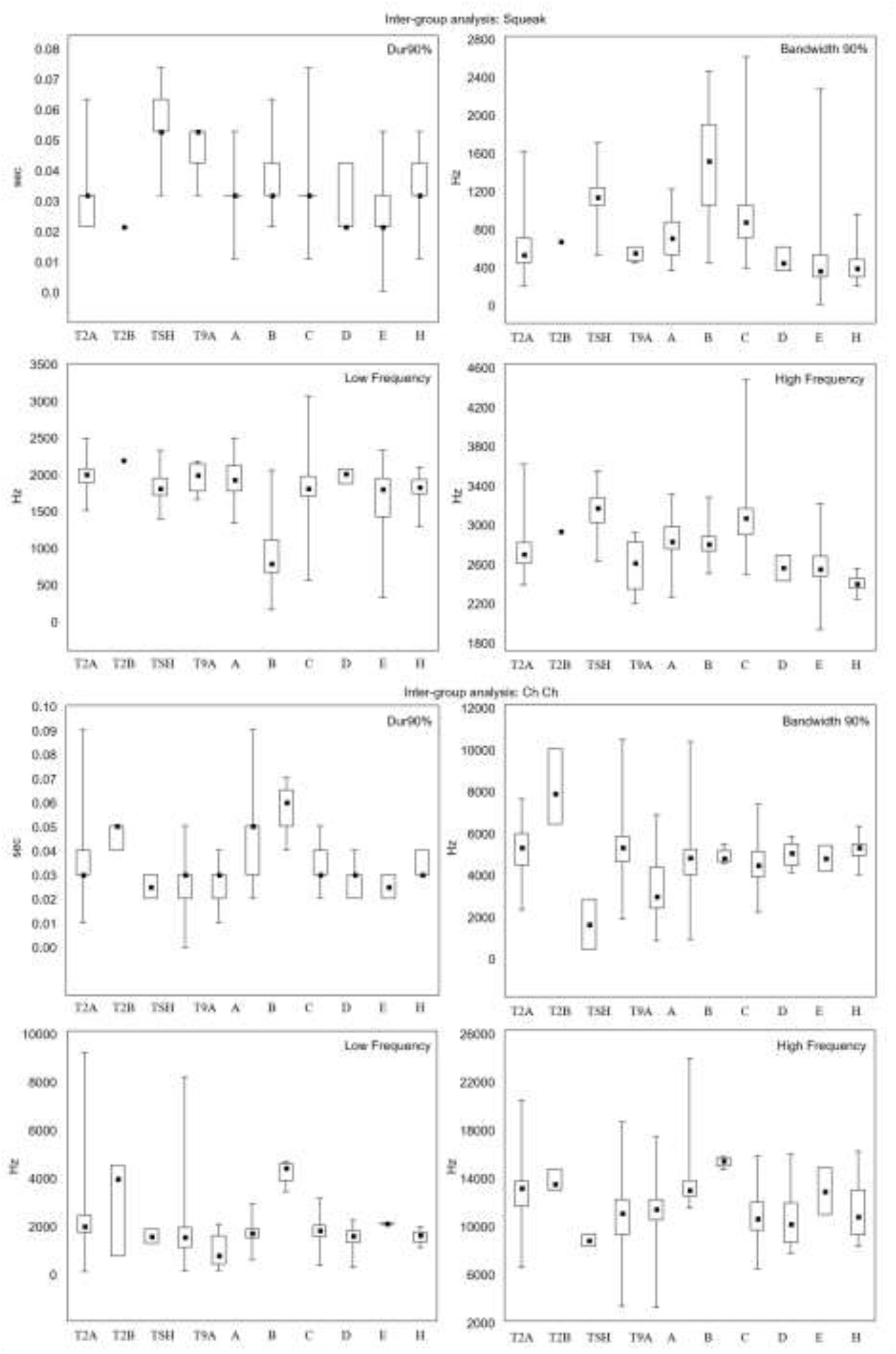
Boxplot plate of four descriptive acoustic measurements (e.g., duration, maximum frequency, minimum frequency, and bandwidth) for the Squeak and Ch Ch across all groups. Median represented by solid box, box plot is 25-75% of call variability, and whiskers are minimum and maximum which do not signify significance, rather distribution of values is depicted. Note that inter quartile ranges are depicted instead of standard error or deviation because of the non-parametric nature. T2A, TSH, and T9A groups are from CREES while A-H are from Villa Carmen. Significant differences illustrated in Table 3

**Fig 4.**
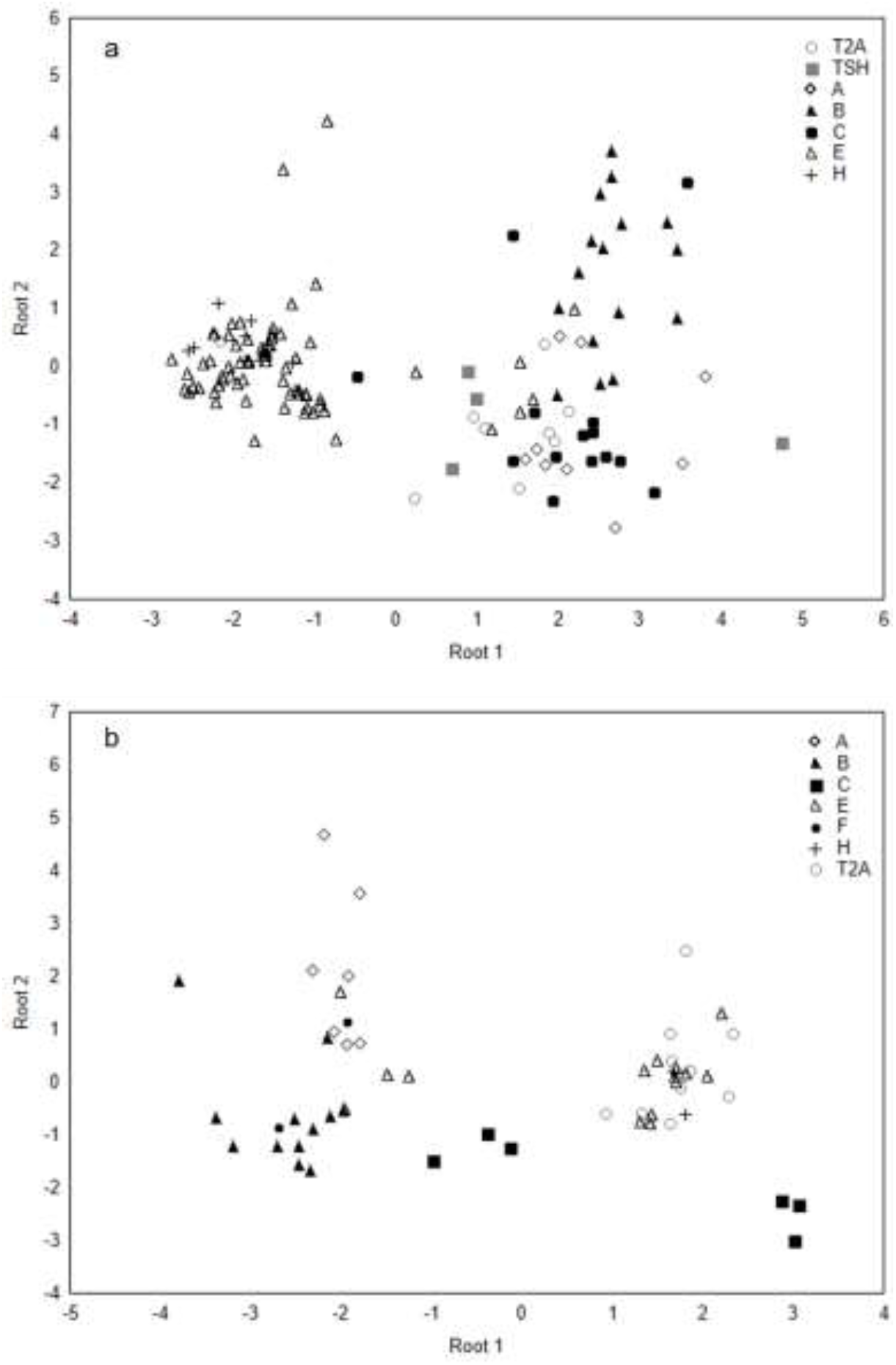
Canonical score scatterplot. The biplot axes are the first two canonical variables. These define the two dimensions that provide maximum separation among groups. We used a 10-fold cross validation in which 90% of the calls were randomly chosen to calculate discriminant functions. Here we present the results of the 10% excluded for testing

Twelve acoustic parameters were significantly different between Villa Carmen and CREES biological field stations (Table 6). Discriminant function analysis identified seven Squeak parameters that significantly distinguished locations: low and high frequency, bandwidth, duration (90%), delta time, IQR duration, and max time – the first of which contributed the greatest discriminatory power; and eight Ch Ch parameters significantly distinguished locations: low and high frequencies, bandwidth, energy, peak frequency, Q1 frequency, frequency (5%) and center time– the last of which contributed the greatest discriminatory power. The two sites were found to be significantly different based on Squeak (Wilks’ Lambda=0.77, F(13,1281)=29.09; p<0.0000) and Ch Ch (Wilks’ Lambda=0.40, F(13,526)=61.77; p<0.0000). Classification accuracy was 93.8% for Squeak (5 out of 13 CREES measurements and 116 out of 116 at Villa Carmen), and 100.0% for Ch Ch. Cumulative significant functions were able to account for 47.7% of variance between locations using the Squeak, and 77.7% using the Ch Ch call.

**Table 6.**
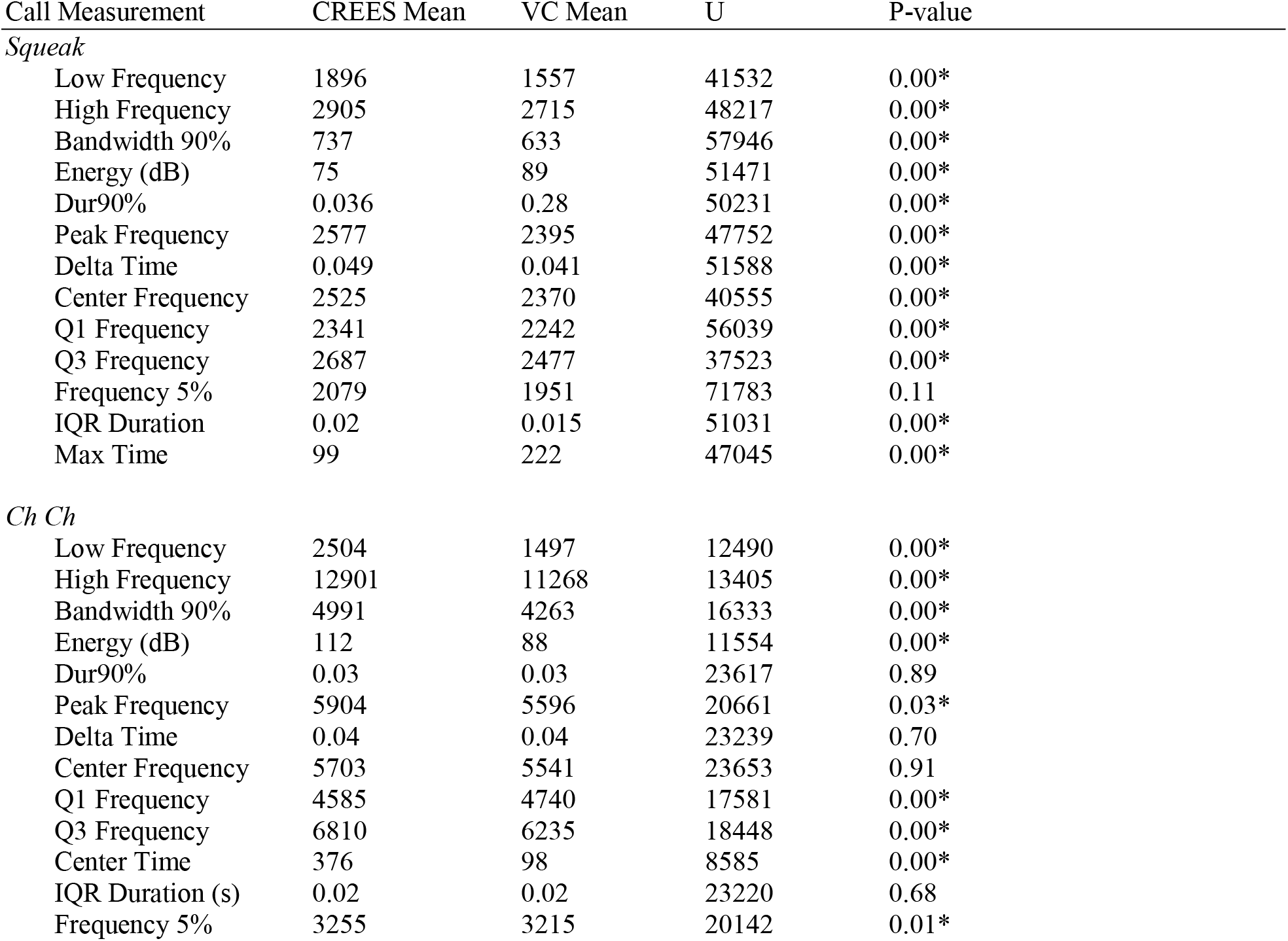
Mann Whitney results comparing group acoustic parameters between distant locations (~10km). Sample size (Ch Ch: CREES 108 and VC 441) and (Squeak: CREES 136 and VC 1150). *All associations significant at p<0.05.

| Call Measurement | CREES Mean | VC Mean | U | P-value |
| --- | --- | --- | --- | --- |
| *Squeak* |  |  |  |  |
| Low Frequency | 1896 | 1557 | 41532 | 0.00* |
| High Frequency | 2905 | 2715 | 48217 | 0.00* |
| Bandwidth 90% | 737 | 633 | 57946 | 0.00* |
| Energy (dB) | 75 | 89 | 51471 | 0.00* |
| Dur90% | 0.036 | 0.28 | 50231 | 0.00* |
| Peak Frequency | 2577 | 2395 | 47752 | 0.00* |
| Delta Time | 0.049 | 0.041 | 51588 | 0.00* |
| Center Frequency | 2525 | 2370 | 40555 | 0.00* |
| Q1 Frequency | 2341 | 2242 | 56039 | 0.00* |
| Q3 Frequency | 2687 | 2477 | 37523 | 0.00* |
| Frequency 5% | 2079 | 1951 | 71783 | 0.11 |
| IQR Duration | 0.02 | 0.015 | 51031 | 0.00* |
| Max Time | 99 | 222 | 47045 | 0.00* |
| *Ch Ch* |  |  |  |  |
| Low Frequency | 2504 | 1497 | 12490 | 0.00* |
| High Frequency | 12901 | 11268 | 13405 | 0.00* |
| Bandwidth 90% | 4991 | 4263 | 16333 | 0.00* |
| Energy (dB) | 112 | 88 | 11554 | 0.00* |
| Dur90% | 0.03 | 0.03 | 23617 | 0.89 |
| Peak Frequency | 5904 | 5596 | 20661 | 0.03* |
| Delta Time | 0.04 | 0.04 | 23239 | 0.70 |
| Center Frequency | 5703 | 5541 | 23653 | 0.91 |
| Q1 Frequency | 4585 | 4740 | 17581 | 0.00* |
| Q3 Frequency | 6810 | 6235 | 18448 | 0.00* |
| Center Time | 376 | 98 | 8585 | 0.00* |
| IQR Duration (s) | 0.02 | 0.02 | 23220 | 0.68 |
| Frequency 5% | 3255 | 3215 | 20142 | 0.01* |

Other differences were observed between groups as well. The Ch Ch was predominately found in a series of two (“in duplicate”) (88.3% of cases); however, four groups also produced calls in triplicate (i.e. Ch Ch Ch). Out of 556 total Ch Ch calls, 65 were in triplicate (11.7%), with 2.8% in T2A, 1.3% in A, 50.4% in B, and 2.8% in E. The distribution of triplicate calls across groups differed significantly from even distribution across groups, with Group B exhibiting nearly four times as many triplicates as expected (p=0.0000). In addition, two groups were observed using ultrasonic frequencies as part of the Ch Ch call (>20kHz): group C (N=3) at Villa Carmen and T2A (N=1) at CREES.

## Discussion

The majority of acoustic parameters for both calls differed significantly between groups and geographic locations, though single acoustic parameters alone were not sufficient to predict group membership. Variance of acoustic parameters overlapped in nearby groups, making absolute classification difficult. However, there was a consistent pattern whereby calls from the same groups and population tended to cluster together based on similar acoustic measurements. Population level classification was more accurate, largely driven by acoustic parameters of the Ch Ch call. Quantitative analysis of acoustic traits may therefore be useful in elucidating group and population level differences and may provide useful insight into the underlying phylogenetic relationships between groups, populations and potentially species of *Aotus*. However, additional recordings are needed both at the group and population levels, preferably with more distant populations included.

We were unable to investigate individual acoustic variability because of our inability to pair calls to specific individuals in a complex environment at night. Based on various other primate studies it is likely that individuals can be differentiated based on vocal signatures (Table 1). However, confirmation of vocal individuality will require either analysis in captivity or pairing video and audio recordings in wild nesting groups. If recordings can be attributed to specific individuals, then acoustic analysis could be used to establish whether individual conspecifics vary predictably in their vocalizations. Establishing the ability to vocally differentiate individuals would be particularly useful for a nocturnal species such as the black-headed night monkey, allowing researchers to study group composition solely based on vocal recordings.

Aside from differences in acoustic parameters, two other acoustic differences were discovered among groups. First, a triplet Ch Ch call was found in recordings from groups T2A, A, B, E. Though relatively rare within the sampled populations (11.7% of cases), over half of these cases were found in Group B. The other groups at Villa Carmen that used the triplet call are likely of the same population since they are isolated on all but one side and in relative proximity to group B (<1300m at furthest extent). The prevalence of the triplet call in Group B suggests that this is not an aberration but rather a consistent modification of a common call. The fact that the triplet call only occurred in certain groups could reflect any number of possibilities including increased prevalence of a particular behavioral context, or a vocal innovation (genetic or learned). Alternatively, it is possible that the presence of both duplicate and triplet Ch Ch calls is the ancestral state and the absence of the triplet call is derived. Either way, additional sampling of nearby groups – coupled with underlying population genetics analysis - would confirm whether this is a relatively unique acoustic irregularity which is independent of underlying population structure, or whether this call variation routinely arises and is widespread. Likewise, being able to obtain calls specific to individuals through video and audio pairing in nests would allow us to decipher whether all individuals within a particular group use the triplet call.

It is uncertain whether the use of ultrasonic frequencies in night monkeys is rare or whether this is a common response to environmental pressures such as inter-species competition for lower frequencies or predator avoidance. Of course, other nocturnal primates (i.e., *Tarsius*, *Galago*, *Microcebus*, *Nycticebus*) and some diurnal neotropical primates (i.e., *Callithrix* and *Cebuella*) produce calls containing ultrasonic frequencies, though only the tarsiers produce calls entirely within the ultrasonic range, with the other species always producing dominant frequencies in the human audible range (Ramsier et al. 2012). In several species, the use of ultrasound appears to be context specific, often in the presence of predators, including humans (e.g. Rahlfs and Fichtel 2010; Gursky-Doyen 2013).

*Aotus* currently consists of eleven described species based on both phenotypic and genotypic evidence. Night monkey taxonomy has been revised considerably based on differences in karyotypes, morphology, molecular sequencing, malaria sensitivity, immunological responses, and geographic isolation (Menezes et al. 2010). Despite this, few specimens from any one study have come from *Aotus nigriceps* despite this species having one of the largest ranges of any *Aotus* species. Moreover, the current taxonomic classification lumps *A. nigriceps* populations from areas with considerably different elevations and from areas separated by significant river systems. Thus, the possibility remains that further evolutionary and conservation management units may exist. Considering the distinct differences in call types previously described between *Aotus* species and the use of quantitative acoustic sampling to differentiate many other primate species, we anticipate that further analysis would prove useful in differentiating population-level or species-level taxonomy.

Finally, *Aotus nigriceps* likely produce more than the three described call types since captive *Aotus* species have exhibited larger vocal repertoires. In captive situations it is easier to record night monkeys at close distances and calls can be induced in different situations, which could facilitate observation of a wider variety of call types. We anticipate that with continued sampling these additional call types could also be recorded in the wild.

## Acknowledgements

We would like to thank the Amazonian Conservation Association (ACA), Villa Carmen Biological Station, and Manu Learning Center (CREES) staff for hosting us, clearing trails, and providing valuable insight into location and behavior of groups. We are indebted to students and staff from the School for Field Studies who assisted with data collection and logistics. We carried out data collection in accordance with the legal requirements of Peru, and with permission of the Amazon Conservation Association and CREES.

